# Pneumococcus drives STAT3 activation of lower airway epithelium in a strain and burden-dependent manner

**DOI:** 10.64898/2026.02.13.705726

**Authors:** Anna Both, Christine Chevalier, Michael G. Connor, Melanie A. Hamon

## Abstract

Crosstalk between respiratory bacteria and human airway epithelial cells initiates cellular immunity, yet the molecular determinants of asymptomatic carriage versus inflammation remain incompletely understood. A key regulator of airway epithelial cellular immunity is the Signal Transducer and Activator of Transcription (STAT3). Once activated by phosphorylation this host transcription factor signals through two cascades, canonical by phosphorylation at tyrosine 705 and non-canonical by phosphorylation at serine 727, which contributes to bacterial clearance and immune defense in pneumonia. Despite this, its role in epithelial responses to *Streptococcus pneumoniae* has not been defined. Revealing these processes are particularly important for understanding the variability in pathogenic potential for the pathobiont *Streptococcus pneumoniae* which triggers either host commensal-like carriage or susceptibility at this interface. Here we investigated the role of STAT3 signaling during pneumococcal challenge using lower airway epithelial cells and two pneumococcal isolates favoring either commensal-like carriage, 6B ST90, or with pathogenic potential, TIGR4. We show the invasive TIGR4 strain drives canonical STAT3 phosphorylation and suppresses non-canonical STAT3 in a burden–dependent manner. This contrasts with the 6B ST90 isolate which requires a 20-fold increase in burden to elicit minimal STAT3 responses. TIGR4 activation correlated with SOCS3 expression, while knockdown of endogenous STAT3 did not alter pneumococcal adherence or epithelial membrane integrity. Mechanistically, STAT3 activation required live bacteria and pneumolysin production but was independent of bacterial hydrogen peroxide. Altogether we reveal previously unrecognized divergence in STAT3 signaling during pneumococcal infection.

## Introduction

The airway epithelium is the first line of defense of the respiratory tract, shielding against inhaled pathogens while simultaneously sustaining a diverse community of commensal microbes. For this, airway epithelial cells balance tolerogenic processes to maintain commensals with pro-inflammatory responses required to eliminate invasive microbes. Maintaining this balance is essential to prevent uncontrolled inflammation which may cause loss of commensals and tissue damage. To date, the crosstalk between bacteria and the host response, which results in either asymptomatic carriage or symptomatic infection is an active area of research for many bacterial species, including *Streptococcus pneumoniae* ^1–6^. An increasing body of evidence shows that airway epithelial cells play an active role in orchestrating the immune response to bacterial challenge rather than serving solely as a structural barrier. Determining how these cells integrate microbial signals to tilt responses toward protective inflammation or maintenance of epithelial–microbial homeostasis is critical. Understanding this will clarify the transition from homeostasis to infection.

An important signal that integrates environmental cues to regulate inflammation, cell survival, and differentiation is the Signal Transducer and Activator of Transcription, STAT3 ^7–9^. The pleiotropic functional effects of STAT3 are mediated through multiple upstream signals engaging cytokine, growth factor, or G protein–coupled receptors. Since most of these receptors (with the exception of receptor tyrosine kinases) lack intrinsic kinase activity, they rely on recruited kinases such as JAKs or SRC to phosphorylate tyrosine 705 (p-STAT3 [Y705]), which leads to STAT3 dimerization, nuclear translocation and finally STAT3-dependent transcriptional program activation ^8^. In the lung, activation is most prominently driven by IL-6 family cytokine receptors via the gp130 subunit (IL-6R, LIFR), while EGFR signaling also contributes critically to epithelial repair ^10,11^. Further modifications to STAT3 exist besides its canonical phosphorylation site at Y705, most prominently, phosphorylation at S727 (p-STAT3 [S727]). Similarly to canonical Y705 phosphorylation, p-STAT3 (S727) acts in the nucleus as a transcription factor^12^. In addition, non-canonical phosphorylation has been shown to have a role in mitochondria and the endoplasmic reticulum ^13–15^.

STAT3 activation in lung epithelial cells has been observed following infection with bacterial pathogens such as *Escherichia coli, Staphylococcus aureus*, *Pseudomonas aeruginosa*, and in response to bacterial products such as the M1 surface protein from *Streptococcus pyogenes* ^10,16–19^. Many studies have documented a direct link between STAT3 activation and neutrophil recruitment, which in turn is necessary to clear bacterial infection. For instance, in a mouse model of *E. coli* pneumonia, it was shown that mice with a conditional homozygotic knock-out of STAT3 in alveolar epithelial cells had defects in neutrophil recruitment, delayed bacterial clearance and increased lung damage compared to controls with functional epithelial STAT3 ^10^. Similarly, in a mouse model of *S. aureus* pneumonia (MRSA, USA300), STAT3 inhibition or deletion in airway epithelial cells led to defective bacterial clearance and increased lung damage, which was due to the STAT3-dependent expression of Reg3γ a protein with bactericidal activity ^16^. Additionally, a chemical STAT3 inhibitor was shown to reduce neutrophil infiltration into the lungs and decrease overall lung inflammation in a mouse prior to challenge with the M1 surface protein of *Streptococcus pyogenes*, a virulence factor commonly associated with toxic shock syndrome ^17^. In another study, STAT3 loss-of-function mutations—as seen in the rare genetic disorder Job syndrome—were analyzed for their impact on the respiratory epithelium, where these mutations contribute to recurrent lung infections and frequent structural lung damage, such as emphysema. These STAT3 mutations led to important defects in epithelial differentiation, including reduced mucocilliary clearance. Furthermore, upon challenge of these cells with *P. aeruginosa*, they observed greatly reduced neutrophil transepithelial migration and impaired bacterial killing ^20^. In line with this, inducible antimicrobial resistance against an otherwise lethal *P. aeruginosa* challenge was achieved through pre-activation of STAT3, through mechanisms that remain unclear ^21,22^. While overall less is known about the function of phosphorylation of S727 compared to Y705 in infection, in the context of host-pathogen interaction, *Helicobacter pylori* infection has been shown to induce p-STAT3 (S727) recruitment to mitochondria in gastric epithelium, most likely regulating autophagy of damaged mitochondria to influence the outcome of bacterial infection ^23^.

Although recent studies have shed light on the role of STAT3 during lung infection, its function in regulating host responses to *Streptococcus pneumoniae* remains understudied. Similarly to other infections, STAT3 activation seems to have a protective role against *S. pneumoniae* infection, mainly through regulation of the antibacterial peptide LL-37 ^24^. However, no studies have focused on the role of STAT3 in the respiratory epithelium, where *S. pneumoniae* typically colonizes, but can also exploit host susceptibility to become invasive leading to pneumonia, sepsis, or meningitis. Indeed, Pneumococcus displays striking intraspecies genotypic and phenotypic heterogeneity in capsule composition and in the expression of virulence-associated factors such as the cholesterol-dependent pore-forming toxin pneumolysin (Ply), H_2_O_2_, and neuraminidases ^25–29^. While certain strains exhibit substantial pathogenic potential, others are predominantly restricted to the commensal niche ^1^. Given this heterogeneity and the distinct pathogenic potential of pneumococcal strains, it is critical to understand how the epithelium integrates microbial signals to balance antimicrobial defense with tissue resilience. We hypothesize that epithelial STAT3 functions as a regulatory node that shapes colonization-permissive and pro-inflammatory responses depending on pneumococcal strain virulence and bacterial load. By analyzing lower airway epithelial cells exposed to pneumococcal strains with differing pathogenic potential, we aim to determine how STAT3 activation coordinates downstream programs of antimicrobial resistance and tissue resilience, providing further insight into the epithelial contribution to host defense.

## Results

### TIGR4 drives STAT3 Y705 phosphorylation and suppresses S727 phosphorylation

STAT3 is a master regulator of immune response, cell survival and differentiation ^30^. To determine whether pneumococcus activated STAT3 in lower airway epithelial cells, we challenged the A549 cell line (type II pneumocytes) for 2h with multiple MOI (5 to 100) of either the invasive Spn TIGR4 (serotype 4) isolate, or the commensal-like 6B (serotype 6B, ST90, clonal complex [CC] 156, lineage F) isolate ^31^. After challenge samples were methanol fixed for immunofluorescence microscopy to quantify nuclear phosphorylated STAT3 tyrosine 705, for canonical activation, or phosphorylated STAT3 serine 727, for non-canonical activation. For this we quantified p-STAT3 (Y705) or p-STAT3 (S727) nuclear signal intensities as fold-change to uninfected within segmented nuclei using DAPI nuclear stain.

Upon TIGR4 challenge at MOI 5, 20, and 100 a clear and significant increase in p-STAT3 (Y705) within the nucleus occurred in comparison to both uninfected and 6B ST90 challenges (Fig. 1A & B). In fact, 6B ST90 challenge did not result in nuclear p-STAT3 (Y705) accumulation until MOI 100 in comparison to uninfected. Even at this high burden the amount of nuclear p-STAT3 (Y705) remained significantly lower than that of TIGR4 at the same MOI (Fig. 1A & B). Stimulation with IL-6 cytokine, a positive control for canonical STAT3 activation for 15 minutes ^32^, showed a significant increase in nuclear p-STAT3 Y705, confirming the responsiveness of STAT3 in the A549 cell line (Sup Fig. 1A). Together, these data demonstrate that the invasive favoring TIGR4 isolate drives canonical STAT3 signaling in A549 epithelial cells across all burdens, as opposed to the commensal-like 6B ST90 strain which requires a 20-fold increase in burden to elicit a response.

**Figure 1:**
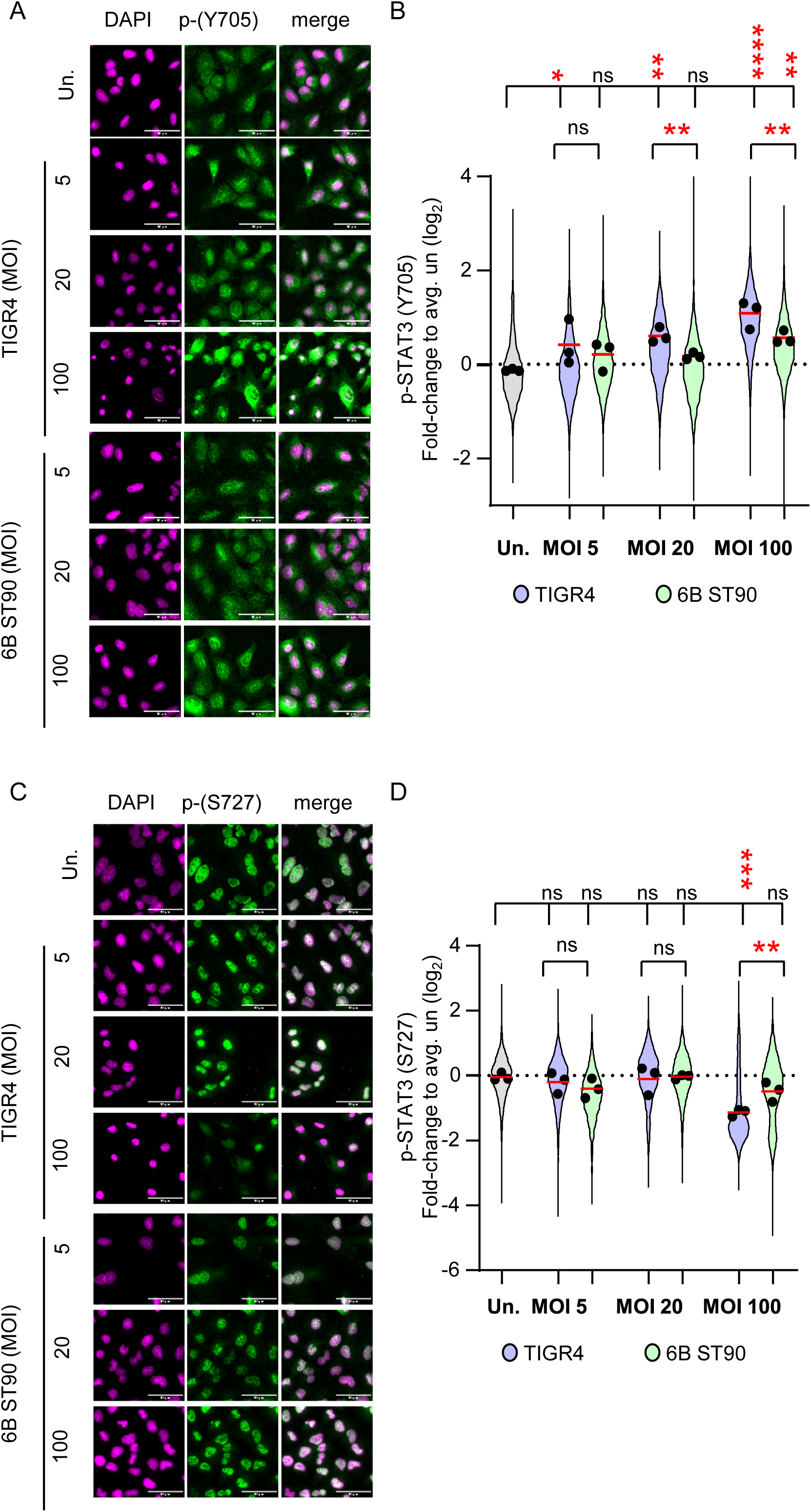
Differential canonical and non-canonical STAT3 Activation in A549 Cells Upon Spn Challenge Is Dependent on Strain and Infection Dose **A**. Representative immunofluorescence images showing nuclear localization of phosphorylated STAT3 at tyrosine 705 (p-STAT3 [Y705]) in A549 cells 2 h after challenge with *Streptococcus pneumoniae* (Spn) strains TIGR4 or 6B at multiplicities of infection (MOIs) of 5, 20, or 100, compared with uninfected controls (un.). Cells were stained with an anti-p-STAT3 (Y705) antibody (green; GFP-labeled secondary antibody) and DAPI for nuclear staining (magenta; pseudocolored). **B.** Quantification of nuclear p-STAT3 (Y705) immunofluorescence (IF) intensity in A549 cells 2 h after infection with Spn TIGR4 or Spn 6B at the indicated MOIs. Nuclear segmentation was based on DAPI staining. IF intensity values were normalized to the mean of the uninfected (un.) control within each experiment and log₂-transformed. Each point represents an individual nucleus (un., grey; TIGR4, purple; 6B, green). Black circles indicate the mean of each of the n = 3 independent experiments, and red horizontal lines denote replicate means. Statistical significance was assessed using two-way ANOVA followed by Tukey’s post-hoc test; ns, not significant (p > 0.05); **p < 0.01; ****p < 0.0001. **C.** Representative immunofluorescence images showing nuclear localization of phosphorylated STAT3 at serine 727 (p-STAT3 [S727]) in A549 cells 2 h after challenge with Spn strains TIGR4 or 6B at MOIs of 5, 20, or 100, compared with uninfected controls (un.). Cells were stained with an anti-p-STAT3 (S727) antibody (green; GFP-labeled secondary antibody) and DAPI for nuclear staining (magenta; pseudocolored). **D.** Quantification of nuclear p-STAT3 (S727) immunofluorescence (IF) intensity in A549 cells 2 h after infection with Spn TIGR4 or Spn 6B at the indicated MOIs. Nuclear segmentation was based on DAPI staining. IF intensity values were normalized to the mean of the uninfected (un.) control within each experiment and log₂-transformed. Each point represents an individual nucleus (un., grey; TIGR4, purple; 6B, green). Black circles indicate the mean of each of n = 3 independent experiments, and red horizontal lines denote replicate means. Statistical significance was assessed using two-way ANOVA followed by Tukey’s post-hoc test; ns, not significant (p≥0.05); *p < 0.05; **p < 0.01; ***p < 0.001; ****p < 0.0001.

Following analysis of canonical STAT3 activation, we evaluated if non-canonical STAT3 signaling, characterized by phosphorylation at serine 727 (S727), was altered upon pneumococcal challenge. Nuclear p-STAT3 (S727) intensity remained largely unchanged at low MOIs for both strains. However, TIGR4 elicited at an MOI 100 a significant decrease in nuclear p-STAT3 (S727) signal intensity (Fig. 1C & D). Finally, by probing whole cell lysates collected 2 h post-challenge we demonstrated total cellular STAT3 was not altered by pneumococcal challenge (Sup Fig. 1B). Overall, these results show that invasive favoring TIGR4 strain activates the canonical STAT3 while suppressing the non-canonical pathway in a bacterial burden dependent manner.

### Pneumococcus alters Expression of STAT3-regulated genes

As TIGR4 challenge increased p-STAT3 (Y705) we tested whether this correlated with downstream STAT3 dependent transcription. A key transcriptional target of STAT3 is *SOCS3* (suppressor of cytokine response 3), which functions to mediate a negative feedback loop to dampen STAT3 activation ^33^. Thus, we assessed whether pneumococcal challenge induced *SOCS3* gene transcription using total RNA collected from A549 cells 2 h post challenge with either TIGR4, 6B ST90, or after a 30-minute stimulation with IL-6 cytokine (50 ng/mL; positive control) for RT-qPCR.

Stimulation with the cytokine IL-6 resulted in 1.5 log_2_ increase in *SOCS3* expression in comparison to uninfected cells after 30 minutes (Fig. 2A). In parallel we assessed whether A549 cells are responsive to IL-6 stimulation by determining *SOCS3* induction 30- and 120-minutes post-stimulation. Indeed, *SOCS3* expression followed a conventional acute stimulus expression curve (Sup. Fig. 2A). TIGR4 challenge at both MOI 20 and 100 induced SOCS*3* transcription, only at a challenge MOI of 100 did 6B ST90 induce *SOCS3* expression in comparison to uninfected (Fig. 2A). Moreover, comparing *SOCS3* expression between TIGR4 and 6B ST90 MOIs showed significant statistical difference (p≤0.05) at MOI 20, but no statistical difference at MOI 100 even though *SOCS3* levels for 6B ST90 remained slightly lower than that of TIGR4 (Fig. 2A). These results aligned with our microscopy analysis of nuclear p-STAT3 (Y705), indicating differential STAT3 activation by pneumococcus leads to SOCS3 expression in a strain- and MOI-dependent manner.

**Figure 2:**
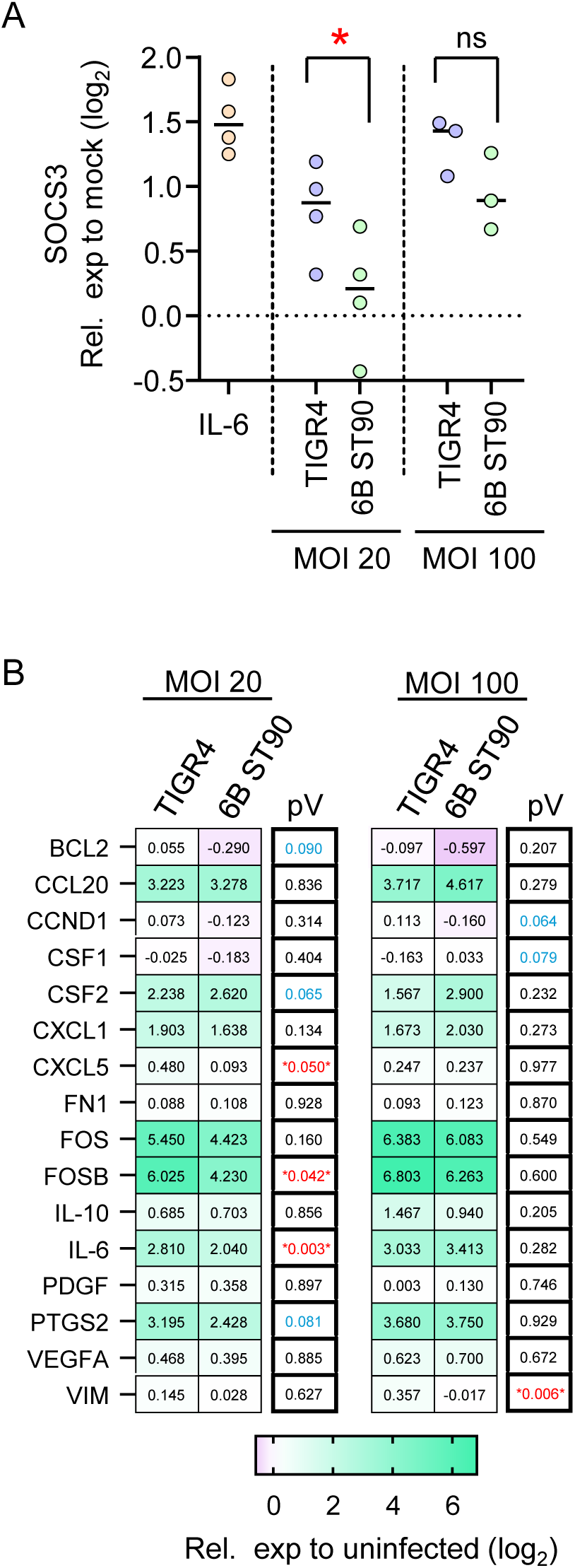
*S. pneumoniae* Infection Drives Expression of STAT3-Regulated Genes in A549 cells **A** Semi-quantitative real-time PCR analysis of SOCS3, a negative feedback regulator of STAT3, in A549 cells following infection with Spn TIGR4 or Spn 6B at MOI 20 or 100 for 2 h. Data are shown as log₂ fold changes relative to the uninfected (Un.) control from n = 4 biological replicates (MOI 20) and n = 3 biological replicates (MOI 100). Gene expression was normalized to the housekeeping gene GAPDH and subsequently to the uninfected control. Statistical comparisons between Spn TIGR4 and Spn 6B at each MOI were performed on −ΔΔCt values using Student’s *t*-test. *p < 0.05; *p < 0.01. **B** Heatmap of real-time PCR results showing expression of selected STAT3-associated genes of interest (GOIs) in A549 cells after 2h-challenge with Spn TIGR4 or 6B at MOI 20 and 100. Plotted values represent log₂ fold changes relative to uninfected. Statistical comparisons were made between Spn strains TIGR4 and 6B for MOI 20 and 100 using Student’s t-test on ΔCt-values. p-values (pV) are stated in the right most column,* any p < 0.05. Color scale reflects relative expression levels (see scale bar).

To further assess the transcriptional output of STAT3 signaling, we profiled the expression of a selection of additional genes reported to be regulated by STAT3 and involved in immune responses or epithelial remodeling. For this we constructed a RT-qPCR panel targeting the STAT3-regulated genes using Harmonizome 3.0 ^34,35^, which aggregates ChIP-seq and ENCODE datasets: BCL2, *CCL20*, *CCND1*, *CFS1, CSF2*, *CXCL1*, *CXCL5*, *FN1*, *FOS*, *FOSB*, *IL-10, IL-6, PDGF, PTGS2, VEGFA* and *VIM*. Total RNA from A549 cells 2 h post-challenge with either TIGR4 or 6B ST90 at MOIs 20 and 100 was used to assess the expression of these targets against unchallenged controls.

Our results showed at MOI 20 TIGR4 significantly increased expression of *CXCL5, FOSB* and *IL-6* in comparison to 6B ST90 at the same MOI (p-value in figure), along with increased expression *BCL2, CSF2* and *PTGS2* which border significance (Fig. 2B). However, at MOI 100 these targets were no longer significant between TIGR4 and 6B ST90 (Fig. 2B). Additionally, stimulation with IL-6 cytokine (50 ng/mL) at 30 and 120 minutes showed a peak elevation for many these targets at 30 minutes, which further substantiates that canonical IL-6 driven STAT3 signaling and transcription is functional in A549 cells (Sup. Fig. 2B). Moreover, at MOI 20 and 100 TIGR4 increased expression over 2-fold (log_2_) of *CCL20, CSF2*, *CXCL1, FOS, FOSB, IL-6, & PTGS2* in comparison to unchallenged cells. Intriguingly, 6B ST90 at either MOI 20 or 100 did alter expression of several of these STAT3 targets (*CCL20, CSF1, CSF2*, *CXCL1, FOS, FOSB, IL-6 & PTGS2)* over unchallenged cells even at an MOI 20 (Sup. Table 1). This suggests that while 6B ST90 does not significantly induce p-STAT3 (Y705), it does engage STAT3 dependent gene transcription, likely though additional signaling cascades. Altogether these data demonstrate that TIGR4 challenge drives a transcriptional program correlated with canonical p-STAT3 (Y705) activation in A549 at lower bacterial burdens in comparison to 6B ST90.

### STAT3 is dispensable for TIGR4 adhesion and epithelial membrane integrity

Previous studies have shown STAT3 to affect bacterial adherence and epithelial cell membrane integrity ^20^. To investigate whether STAT3 altered *S. pneumoniae* adherence or host membrane integrity during challenge, we used RNAi mediated knockdown of STAT3. We first assessed STAT3 knockdown efficiency in A549 airway epithelial cells using single siRNA for 48 h by immunofluorescence microscopy. This resulted in greater than 50% decrease in nuclear STAT3 quantities regardless of bacterial challenge or IL-6 stimulation in comparison to scrambled (scr) negative control (Sup. Fig. 3A & B). To determine pneumococcal adherence, STAT3 RNAi and scr treated A549 cells were washed to remove non-cell associated bacteria before collecting cell associated bacteria 2 h post-challenge with either TIGR4 or 6B at MOIs 20 and 100. These samples were serially diluted to determine bacterial burdens by CFU counts under the RNAi conditions. Knockdown of STAT3 had no impact on either TIGR4 or 6B cell-associated bacteria at either MOI (Fig. 3A).

**Figure 3:**
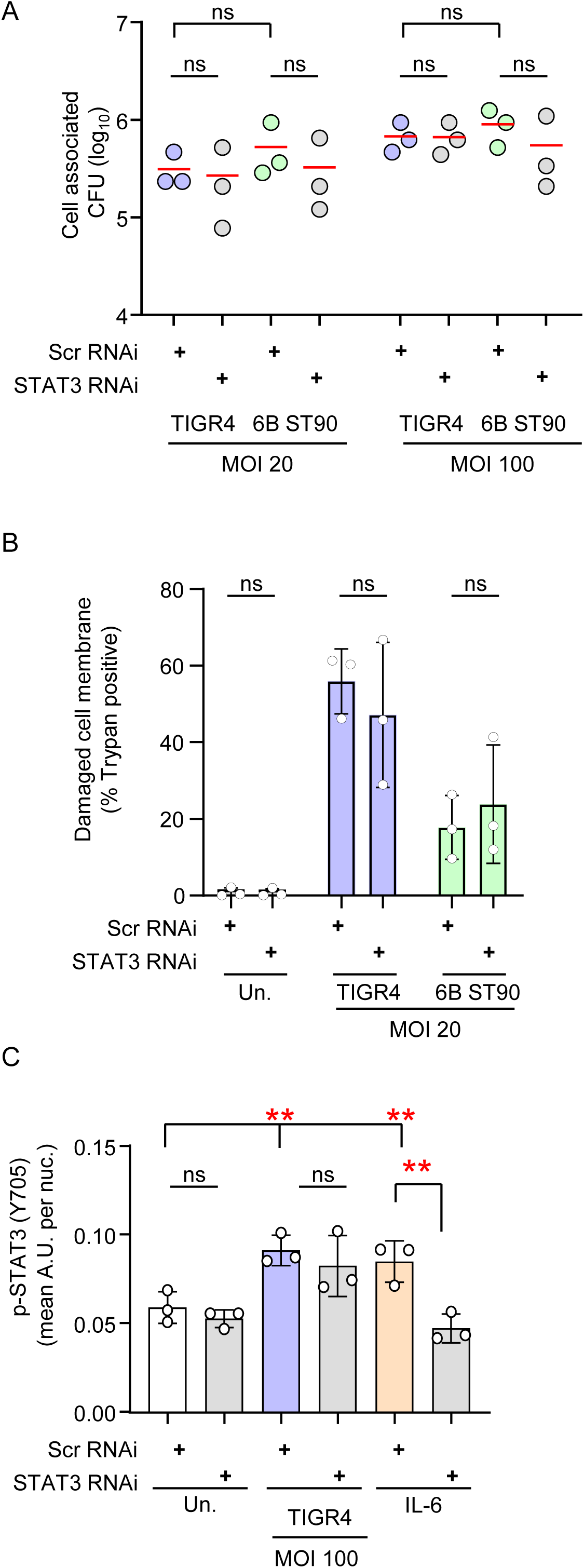
STAT3 knockdown does not affect A549 cell integrity during *S. pneumoniae* challenge **A**. Adhesion of Spn to A549 cells after 48 h of transfection with STAT3-targeting RNAi (STAT3 RNAi) or scrambled control RNAi (scr RNAi). Cells were challenged for 2 h with Spn TIGR4 or Spn 6B at MOIs of 20 or 100, after which adherent bacteria were quantified as CFUs from n = 3 independent biological replicates. **B**. Quantification of trypan blue–positive nuclei in RNAi-treated A549 cells following a 2 h challenge with Spn TIGR4 or Spn 6B at MOI 20. Four to six microscopy fields per condition and experiment were manually analyzed, and the mean percentage of trypan blue–positive cells was determined. Percentages from individual biological replicates (n = 3) are shown. Statistical comparisons between STAT3 RNAi and scr RNAi were performed for each condition using a two-sided Student’s t-test. **C**. Quantification of mean nuclear phosphorylated STAT3 (p-STAT3 [Y705]) fluorescence intensity in RNAi-treated A549 cells (n = 3 biological replicates) after 48 h of RNAi treatment, followed by either a 2 h challenge with Spn TIGR4 at MOI 100 or stimulation with IL-6 (50 ng/mL) for 15 min. Nuclear segmentation was based on DAPI staining. Statistical analysis was performed using two-way ANOVA followed by Tukey’s post-hoc test. Statistical significance: ns, not significant (p > 0.05); *p < 0.05; **p < 0.01.

Having established knockdown of STAT3 did not change adherence we tested if reduced levels of STAT3 increased A549 cell susceptibility to pneumococcal induced membrane damage during challenge. We previously showed at an MOI of 20 TIGR4 challenge, as opposed to 6B ST90, reduces plasma membrane integrity using a trypan blue exclusion assay as cells with damaged plasma membranes accumulate the dye ^31,36^. While TIGR4 caused more membrane damage than 6B, there was no significant difference in trypan blue uptake between STAT3 knockdown and the scrambled (scr) control cells (Fig. 3B; representative images Sup. Fig. 3C). Furthermore, we used a lactate dehydrogenase (LDH) release assay to quantify cell death over the 2 h bacterial challenge to corroborate that not all trypan positive cells are dead (Sup. Fig. 3D). Overall, these results show that STAT3 does not impact pneumococcal adherence to A549 cells and it does not influence epithelial plasma membrane integrity or cellular cytotoxicity during bacterial challenge.

### Reduction in total STAT3 does not alter TIGR4 driven nuclear p-STAT3 (Y705) levels

Having established that STAT3 had a dispensable role in TIGR4 adherence and host epithelial cell resilience to challenge, we tested whether RNAi mediated decrease in total STAT3 affected levels of TIGR4 driven p-STAT3 (Y705). For this we challenged A549 cells under RNAi STAT3 or scrambled (scr) conditions for 2 h with either TIGR4 MOI 20 or 100 alongside a positive control stimulated for 15 minutes with IL-6 (50 ng/mL) for immunofluorescence imaging of nuclear p-STAT3 (Y705).

Interestingly there was no difference in the basal level of nuclear p-STAT3 Y705 for uninfected samples when STAT3 was knocked down in comparison to the scrambled control (Fig. 3C), even though knockdown efficiency of total STAT3 was greater than 50% (Sup. Fig. 3B). Similarly, our results clearly showed that under TIGR4 challenge at either MOI, even with comparable reduction in total cellular STAT3 (Fig. 2B), there was minimal impact on nuclear p-STAT3 (Y705) levels in comparison to the scrambled siRNA control (Fig. 3C). However, cells stimulated with IL-6 resulted in significant (p≤0.01) reduction in nuclear p-STAT3 (Y705)( Fig. 3C). Altogether these data suggest that different pathways are driven by TIGR4 and IL-6 to trigger p-STAT3 (Y705), as knockdown of total STAT3 only meaningfully impacted IL-6 induced p-STAT3 (Y705) levels.

### Canonical STAT3 activation requires Live Pneumococcus and Pneumolysin

Having established that STAT3 had a dispensable role in TIGR4 adherence and host epithelial cell resilience to challenge, we investigated whether increased p-STAT3 (Y705) required active bacterial mechanisms or the pneumococcal factors of Pneumolysin toxin (PLY) and hydrogen peroxide, which are encoded by the respected genes *ply* and *spxB* (pyruvate oxidase). Herein, A549 cells were challenged for 2 hrs with either live TIGR4, paraformaldehyde (PFA) inactivated TIGR4, TIGR4Δ*ply* or TIGR4Δ*spxB* at MOI 20 and 100 prior to methanol fixation and immunofluorescence staining for nuclear p-STAT3 (Y705) as a marker for canonical STAT3 activation.

We show in comparison to unchallenged cells that live TIGR4 at either MOI induced nuclear p-STAT3 (Y705) in contrast to PFA inactivated TIGR4 which did not (Fig. 4A). These results indicate that bacterial surface ligands alone are insufficient to activate STAT3 signaling. We next focused on Pneumolysin toxin (Ply), a cholesterol-dependent pore-forming toxin (PMID: 32714314). At either MOI the TIGR4Δ*ply* did not induce STAT3 phosphorylation in contrast to the parental wildtype strain, suggesting that Pneumolysin was essential for this phenotype (Fig. 4B). To test this exclusivity, we exposed A549 cells for 2 hrs to recombinant Pneumolysin of serotype 4 (same as TIGR4) at 0.1, 2 and 10 nM concentrations. However, treatment with purified Pneumolysin at any concentration tested failed to increase nuclear p-STAT3 (Y705) in comparison to untreated cells (negative control) or TIGR4 challenge at MOI 100 (positive control, Sup. Fig. 4D). Thus, Ply is required by live bacteria but is not sufficient in and of itself to drive the canonical STAT3 phenotype. Finally, we tested whether bacterial hydrogen peroxide production was required to drive nuclear increase in p-STAT3 (Y705) by comparing a TIGR4Δ*spxB* to its parental wildtype strain. Our results show challenge at either MOI with TIGR4Δ*spxB* displayed comparable STAT3 phosphorylation as the parental TIGR4 strain (Fig. 4C). These data indicate that hydrogen peroxide production is not needed for p-STAT3 (Y705).

**Figure 4:**
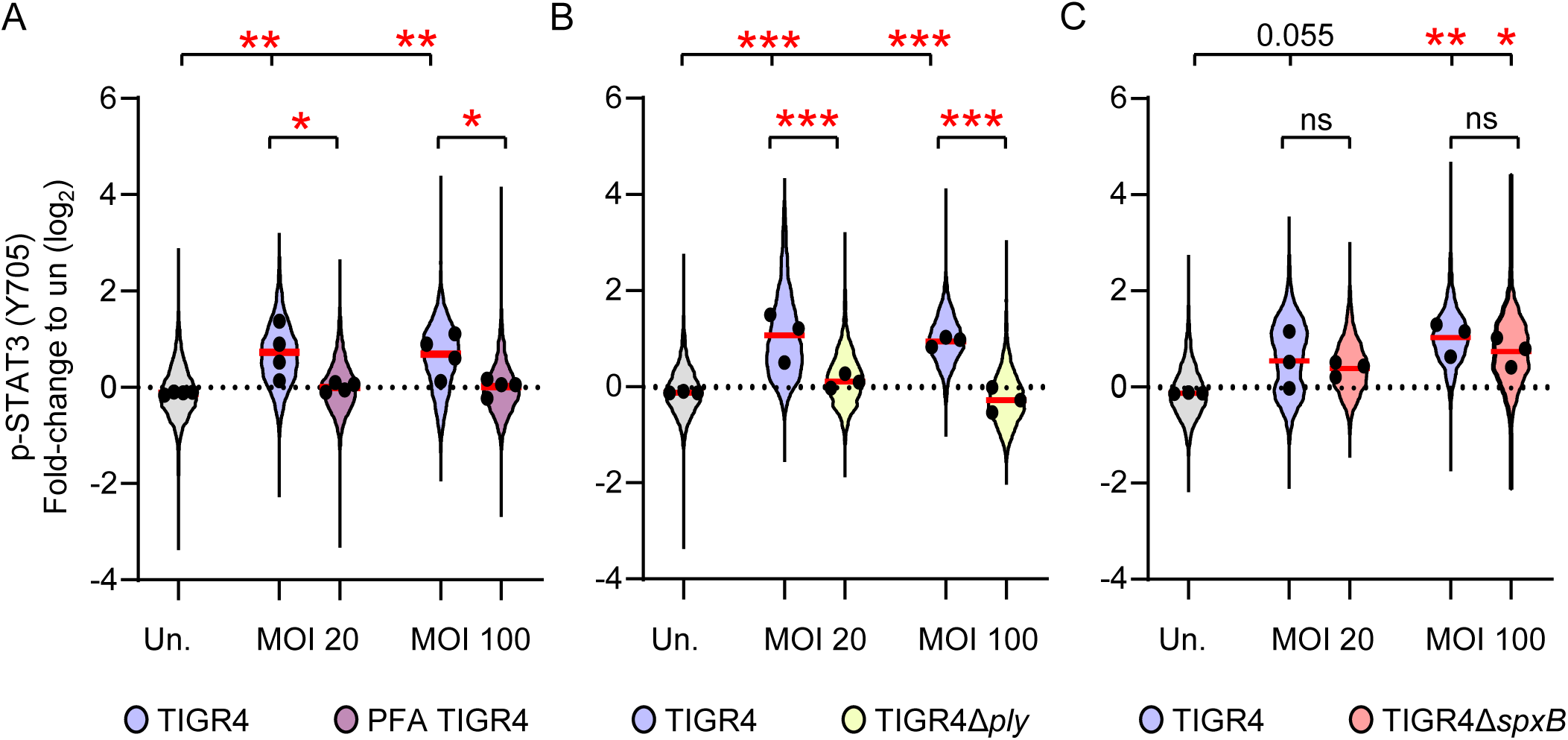
Live *S. pneumoniae* and Pneumolysin Production Are Required to Trigger Canonical STAT3 Activation Quantification of phosphorylated STAT3 immunofluorescence intensity (IF) over the nucleus in A549 cells, segmented based on DAPI nuclear staining. Fluorescence intensities were normalized to the mean of the uninfected control (un.) for each experiment and log₂-transformed. Each colored point represents an individual nucleus; black circles indicate the mean of biological replicates, and horizontal lines show the overall mean. Statistical significance was assessed using two-way ANOVA followed by Tukey’s post-hoc test. *p < 0.05; **p < 0.01; ***p < 0.001. **A.** Quantification of p-STAT3 (Y705) IF intensity over the nucleus of A549 cells challenged with live Spn TIGR4 and PFA-inactivated TIGR4 (PFA TIGR4) at MOIs of 20 and 100. Color coding: un. (grey), live TIGR4 (purple), inactivated TIGR4 (mauve). (n = 4). **B.** Quantification of p-STAT3 (Y705) IF intensity over the nucleus of A549 cells challenged withSpn TIGR4 and Spn TIGR4Δ*ply* (pneumolysin-deficient) at MOIs 20 and 100. Color coding: un. (grey), TIGR4 (purple), TIGR4Δ*ply* (yellow). (n = 3). **C.** Quantification of p-STAT3 (Y705) IF intensity over the nucleus of A549 cells challenged withSpn TIGR4 and Spn TIGR4Δ*spxB* (pyruvate oxidase-deficient) at MOIs 20 and 100. Color coding: un. (grey), TIGR4 (purple), TIGR4Δ*spxB* (salmon). (n = 3).

Likewise, analysis of the TIGR4-dependent non-canonical STAT3 phenotype showed a requirement for live bacteria and Ply expression (Sup. Fig. 4A,B). PFA-inactivated bacteria and TIGR4Δply failed to induce the reduction in nuclear p-STAT3 (S727) observed with the wild type, whereas TIGR4Δ*spxB* showed a similar tendency to the wild type in reducing p-STAT3 (S727) at higher MOI (Sup. Fig. 4C).

Altogether, these results demonstrate that canonical STAT3 activation and non-canonical STAT3 suppression in A549 cells require live TIGR4 bacteria harboring an intact *ply* gene. Furthermore, while bacteria producing Pneumolysin toxin is essential for p-STAT3 (Y705) additional interactors at the host–pathogen interface may contribute as recombinant Pneumolysin toxin alone failed to induce p-STAT3 (Y705).

## Discussion

While the majority of *Streptococcus pneumoniae* (the pneumococcus) isolates are asymptomatically carried within the human nasal passage and upper airways, some are responsible for symptomatic diseases such as bacterial pneumonia ^6^. However, it remains an active area of research to uncover mechanisms at the pneumococcal – host interface explaining variations in host outcomes that contribute to the development of pneumonia. STAT3 is a central mediator in epithelial cell immunity and resilience to insult during pneumonia ^9^. As such, we studied whether pneumococcal strains associated with invasive disease (TIGR4) or commensal-like interaction (6B ST90) activated STAT3 in A549 epithelial cells, which are immortalized type II pneumocytes. We discovered low burdens of TIGR4 drive canonical phosphorylation of STAT3 (Y705) and suppressed non-canonical STAT3 phosphorylation (S727). In contrast, 6B ST90 required 5-20fold higher bacterial burdens to elicit even a marginal STAT3 response. Our investigation further showed live bacteria and pneumolysin toxin (Ply), a pore-forming CDC toxin ^37^, were required for the TIGR4 phenotype. However, functional knock-down of STAT3 did not influence pneumococcal adherence or epithelial cell integrity in A549 cells. Altogether, we demonstrate for the first time pneumococcal activation of STAT3 in airway epithelial cells is strain- and burden-dependent.

The divergence between TIGR4 and 6B ST90 to drive STAT3 phosphorylation at Y705 while reducing the levels of phosphorylated STAT3 (S727) across multiple burdens indicates a pronounced difference in signal transduction. Altogether these data provide strong evidence that TIGR4 induction of STAT3 (Y705) is likely though the crosstalk of additional transduction components or signaling cascades, such as NF-kB. Moreover, reduction in total cellular pool of STAT3, by RNAi mediated knockdown, had little effect on A549 epithelial cell membrane integrity or on TIGR4 driven STAT3 phosphorylation at Y705. Specifically, Y705 levels under RNAi mediated STAT3 knockdown stand in contrast to the observed 40% reduction in Y705 under canonical IL-6 stimulation in the same conditions. Finally, we show challenge of airway epithelial cells with either pneumococcal strain or IL-6 increased *SOCS3* expression, a known STAT3 dependent transcript ^38^.

Contextually signaling through interleukin receptors that contain the subunit glycoprotein 130 (gp130)(e.g. IL6R, LIFR), non-gp130-interleukin receptors (e.g. IL21R), certain growth factor receptors (e.g. EGFR, GM-CSFR), and G-protein-coupled receptors have all been shown to converge on STAT3 activation^12^. Reports show that phosphorylated STAT3 (Y705) and S727 may intrinsically regulate each other with S727 phosphorylation accelerating Y705 dephosphorylation or promoting STAT3 degradation in some cellular contexts^39–41^. To our knowledge, such suppression of S727 phosphorylation during bacterial infection has not been reported. In contrast, viral and *Helicobacter pylori* infection models show increased p-S727, largely in the mitochondrial compartment ^23^. It remains unknown whether TIGR4 induced p-STAT3 (Y705) requires gp130, Src or JAKs, or whether the increase in p-STAT3 (Y705) is the result of a shift in host epithelial response to reinforce antibacterial or metabolic defenses against pneumococcal insult.

During pneumococcal pathogenesis cell adherence and key bacterial products, Pneumolysin toxin (Ply) and hydrogen peroxide encoded by the *ply* and *spxB* genes respectively, are known to shape host epithelial responses. Adherence of TIGR4 and 6B ST90 to airway epithelial cells was not impacted by STAT3. However, TIGR4 driven STAT3 (Y705) phosphorylation was dependent on live bacteria and Ply, but not hydrogen peroxide. In contrast, a recent mouse lung infection model showed that treatment with TLR2/6 and TLR9 agonists mediated epithelial STAT3 activation via reactive oxygen species (ROS)-dependent, ligand-independent activation of epithelial growth factor receptor (EGFR), conferring protection against subsequent *P. aeruginosa* challenge. It is worth noting Pneumolysin toxin alone has been shown to disrupt ion balances and trigger intracellular increases in ROS ^42,43^. However, when we tested recombinant Pneumolysin toxin nuclear levels of phosphorylated STAT3 (Y705) remained unchanged. Further detailed investigations to determine the contribution of Ply, and potentially unknown bacterial proteins, to the Y705 and S727 phenotypes are needed.

STAT3 activation is an early initiator of anti-bacterial responses during lung infection to many bacterial species due to its significant role as a transcription factor driving expression of immune signaling molecules, including several needed for neutrophil recruitment ^10,44–47^. Recently, investigation of STAT3 mutations associated with Job syndrome - a condition characterized by heightened susceptibility to bacterial pneumonia—have linked these mutations to defective STAT3 phosphorylation at Y705 and reduced neutrophil recruitment ^20^. Here multiple mutations, including V637M and Y657S located near the Y705 site, resulted in diminished expression of CXCL2 and IL-6 and recruitment of neutrophils when lung epithelial cells were stimulated with IL-6 or *P. aeruginosa* ^20^. Indeed, a critical aspect of pneumococcus induced pneumonia and host survival is shaped by early neutrophil influx ^48,49^. Perplexingly, our study showed CXCL1, also needed for neutrophils, and IL-6 were induced by both TIGR4 and 6B ST90, however only TIGR4 increased STAT3 (Y705) phosphorylation. Because we, and others, have shown several 6B isolates to favor carriage with a lower risk for pneumococcal disease ^31,50^, an intriguing hypothesis is that pneumococcal burden dependent increases in STAT3 (Y705) may impact bacterial clearance and the onset of pneumonia potentially through how neutrophils are recruited. Therefore, defining the signaling pathways that differentially drive Y705 versus S727 STAT3 phosphorylation may have implications beyond pneumococcus and extend to other respiratory pathogens.

Overall, our study reveals that airway epithelial cell STAT3 response is pneumococcal load and strain-specific. We provide strong evidence that the functional and transcriptional consequences of canonical p-STAT3 (Y705) activation and non-canonical p-STAT3 (S727) suppression are likely to influence pneumococcal disease onset and need further study.

## Materials and Methods

### Cell Culture

Human A549 epithelial cells (ATCC ref# CCL-185) were maintained in F-12K medium (Gibco; #21127030) with 10% fetal calf serum (FCS) and 2 mM L-glutamine (v/v) at 37 °C 5% CO₂. Two days prior to bacterial challenge, cells were seeded in tissue culture 24-well or 6-well plates at 5 × 10⁴ cells/well or 2 × 10⁵ cells/well respectively. For immunofluorescence microscopy, 16 mm glass coverslips were acid-washed, rinsed three times in distilled water, UV-treated for 15 min, and placed in 24-well plates prior to cell seeding.

### *Streptococcus pneumoniae* culture and cell challenge

Bacterial strains (Table S1) were grown statically in brain heart infusion (BHI) broth (Gibco, #237300) at 37 °C 5% CO₂ to mid-exponential phase (OD₆₀₀ = 0.5–0.6). Bacteria were harvested by centrifugation (1,500 xg; 10 min), washed three times in PBS, and diluted to desired multiplicities of infection (MOI 1:1, 5:1, 20:1, 100:1) in either F-12K medium with 10% FCS (for A549). Bacterial challenge studies were synchronized at 200 × g for 5 min at room temperature prior to incubation at 37 °C 5% CO₂ for 2 hr experimental challenge. Where indicated as positive STAT3 control, IL-6 (50 ng/mL) was added to cell culture media 15mins prior to experiment termination for immunofluorescence, and at 30min for qPCR.

For inactivated Spn, a mid-exponential phase culture was harvested as above, then suspended in 4% paraformaldehyde (PFA) for 20 min at room temperature. Inactivated bacteria were washed three times in PBS with a sample plated and grown as above to confirm inactivation. Defined numbers of inactivated bacteria were stored at –20 °C based on CFU to OD conversion factors. These aliquots were used to dilute MOI numbers for challenges. This method preserves bacterial surface integrity, including pathogen-associated molecular patterns (PAMPs)^31,51,52^.

### Adherence assay

Spn challenged cells in 24-well plates were washed twice with PBS prior to host cells lysis with 200 µL distilled water for 15 min at 37 °C, 5% CO₂. Lysates were resuspended by thorough pipetting, serial diluted on 5% horse blood Columbia agar plates and grown at 37 °C, 5% CO₂ overnight to determine bacterial numbers (CFU).

### Immunofluorescence microscopy

Cells grown on coverslips were fixed post-challenge in 100% ice-cold methanol for 6 min at –20 °C, washed three times in PBS and blocked with 5% BSA for 1 h at room temperature. Primary antibody staining was carried out overnight at 4 °C in 5% BSA with the following antibodies (all at 1:200 dilution): Anti-STAT3 (124H6) Mouse mAb (Cell Signaling #9139), Anti-p-STAT3 (Y705) (D3A7) XP Rabbit mAb (Cell Signaling #9145), Anti-p-STAT3 (S727) Rabbit mAb (Abcam #ab32143). Coverslips were washed three times in PBS prior to secondary staining for 1 h at room temperature with Alexa Fluor 488-conjugated goat anti-goat IgG (1:1000; Invitrogen #A11034) and Alexa Fluor 594-conjugated goat anti-mouse IgG (1:1000; Invitrogen #A11005) in 5% BSA/PBS, with 300 nM DAPI (Invitrogen #D1306). Coverslips were washed three times in PBS, once in distilled water, and mounted using ProLong™ Gold Antifade Mountant (Invitrogen #P36930).

### Image acquisition and analysis

Images were acquired on a Cytation 5 microscope (BioTek) with Gen5 Image+ software v3.16. Image processing included background flattening (68 µm rolling ball radius) and deconvolution (PSF auto setting for 20× objective). Quantification was performed in CellProfiler v4.2.6 ^53^ by segmenting on the DAPI nuclear stain (primary objects) with nuclear signal intensities for STAT3, p-STAT3 (Y705), and p-STAT3 (S727) measured. Values were normalized to the uninfected control mean for each experiment, log₂-transformed, and analyzed STAT3 RNAi.

Coverslips in 24-well plates seeded with 1.25 × 10⁴ cells/well 24 hr prior to transfection to achieve ∼30% confluence. Transfection was performed with Lipofectamine RNAiMAX (Thermo Fisher #2762333) as per the manufacturer’s protocol. Briefly, 20 nM STAT3-targeting siRNA (Silencer Select #4390824) or scrambled control (Ambion #AM4611) were mixed with RNAiMAX reagent in Opti-MEM (Thermo Fisher #31985062), incubated for 20 min, and added to antibiotic-free medium. Knockdown proceeded for 48 h before bacterial challenge. STAT3 knockdown was validated by immunofluorescence microscopy of STAT3 and p-STAT3 (Y705).

### Trypan Blue Membrane Integrity Assay

To assess membrane integrity, A549 cells were stained with 0.2% trypan blue in PBS for 10 min at 37 °C post-challenge, washed, and fixed in 4% PFA for 20 min. After three PBS washes, cells were imaged in brightfield using an EVOS FL microscope at 10x magnification (Thermo Fisher). For each condition, 4–6 fields of view (based on variability) were analyzed. Trypan blue-positive (dark nuclei) cells were counted and expressed as a percentage of total cells.

### Lactate dehydrogenase (LDH) Release Assay

Lactate dehydrogenase (LDH) release was measured with the Cytotox 96® Non-Radioactive Cytotoxicity Assay (Promega #G1780). Supernatants collected 2 hr post-infection, centrifuged at 10,000 × g for 5 min to remove bacteria, and mixed 1:1 with reaction reagent. After 30 min incubation in the dark, stop solution was added, and absorbance (490 nm) read on a Cytation 5 plate reader. Absorbance of blank medium was subtracted, and values were normalized to a detergent-induced 100% lysis control.

### Quantitative Real-Time PCR (qPCR)

Total RNA was extracted using the RNeasy Mini Kit (Qiagen #74104). RNA concentrations were determined by a NanoDrop spectrophotometer, and 5 µg were reverse-transcribed into cDNA using SuperScript IV with poly-T primers (Thermo Fisher #18090050 and #18418020). qPCR was performed with iTaq Universal SYBR Green Supermix (Bio-Rad #1725124) on a CFX384 cycler (Bio-Rad), using 2 ng cDNA per 10 µL reaction, run in technical duplicates. Cycling conditions included a denaturation step and a 60 °C annealing/extension step for 30 s over 40 cycles, followed by melt curve analysis. All amplicons showed a single melt peak. Expression levels were normalized to *GAPDH* using the ΔΔCt method ^54^. Primer sequences are listed in Table S1.

### Immunoblotting

Cells were lysed in RIPA buffer (Thermo Fisher Scientific #89900) supplemented with PhosSTOP phosphatase inhibitor cocktail (Roche, #4906845001), Sodium Butyrate (10 mM, Sigma), PMSF (1 mM, Sigma), and cOmplete protease inhibitor cocktail (Roche #11836170001). Laemmli buffer with 5% β-mercaptoethanol (Sigma) was added to the lysates, followed by 5 minutes of sonication and 5 minutes of boiling at 95 °C. Proteins were separated on 4–15% Mini-PROTEAN TGX Stain-Free precast SDS-PAGE gels (Bio-Rad, #4568086) and transferred to PVDF membranes (Bio-Rad #1620177 Transblot Turbo). Membranes were blocked for 1 hour at room temperature in TBST 5% BSA (Sigma). Primary antibodies diluted in TBST 5% BSA: STAT3 (mouse mAb(12H6), Cell Signaling Technology #9139S, 1:1,000), Actin (mouse mAb, (AC-15) Sigma #A5441; 1:10,000), were incubated overnight at 4 °C. Following three washes (5 min each) in TBST, membranes were incubated for 1 hour at room temperature with HRP-conjugated secondary antibody (anti-mouse IgG HRP, Millipore #BI2413C) diluted 1:10,000 in TBST with 5% BSA. Membranes were washed 3 times in TBST prior to chemiluminescent detection (Clarity Western ECL Substrate Bio-Rad #1705060) and imaging on a ChemiDoc Imaging System (Bio-Rad). Densitometric quantification was performed using Image Lab Software (Bio-Rad).

### Statistical analysis

If data were normally distributed (Shapiro–Wilk test), comparisons were performed using Student’s t-test, one-way ANOVA, or two-way ANOVA with appropriate corrections (Dunnett’s or Tukey’s). If not, Kruskal–Wallis non-parametric testing was used. All analysis was performed using GraphPad Prism software.

## Data Availability

All data in the present study is available upon request from the corresponding authors.

## Code Availability

No custom code or software was used in the manuscript.

## Acknowledgements

We thank Emmanuelle Varon and Thomas Kohler for the *S. pneumonie* strains used. AB is supported by a Walter Benjamin Fellowship given by Deutsche Forschungsgemeinschaft (DFG, German Research Foundation, grant number 469972649) and the French Government’s Investissement d’Avenir program, the Laboratoire d’Excellence ‘‘Integrative Biology of Emerging Infectious Diseases”. MGC is supported by a Springboard to Independence grant (AirwayStasis) from the French Government’s Investissement d’Avenir program, the Laboratoire d’Excellence ‘‘Integrative Biology of Emerging Infectious Diseases” (ANR-10-LABX-62-IBEID) and an Agence nationale de la recherche Jeunes Chercheuses et Jeunes Chercheurs (JCJC) “CommensalNose” (ANR 24 CE15 7975 01). This project was supported by the lnstitut Pasteur. the Fondation iXCore-iXLife, the EMBO Young Investigator Program, the Agence Nationale de la Recherche (ANR-23-CHBS-0001 ChromaBac). M.A.H. is a member of the Laboratoire d’Excellence “Integrative Biology of Emerging Infectious Diseases” Agence Nationale de la Recherche (ANR-10-LABX-62-EIBID).

## Author contributions

Conceived and designed all experiments: AB, MGC and MAH. Performed and analyzed data for all experiments: AB & MGC with specific contributions from CC (western blot). AB wrote the original manuscript, MGC and MAH edited and reviewed the final version. MGC and MAH supervised the research. All authors approved the final manuscript.

## Conflict of interest statement

The authors declare no conflict of interest.

---

**Supplementary Figure S1:**
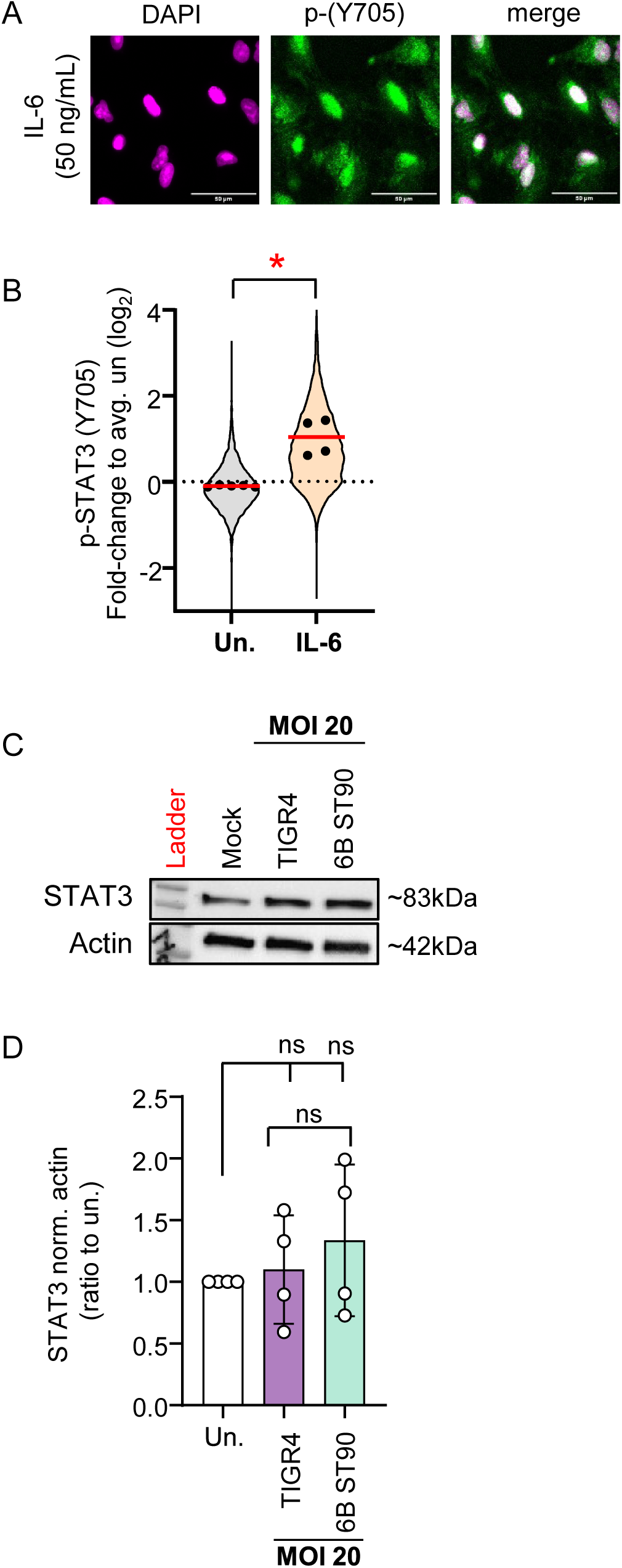
Total STAT3 Protein Levels, Nuclear Size Independence, and Cell Integrity in A549 Cells During *S. pneumoniae* Challenge **A** Representative immunofluorescence images showing nuclear localization of phosphorylated STAT3 (p-STAT3 [Y705]) in A549 cells 15 minutes after stimulation with IL-6 (50ng/mL). Cells were stained with a primary anti-p-STAT3 (Y705) antibody (green; GFP-labelled secondary) and DAPI for nuclear staining (magenta; pseudocolored). **B** Quantification of nuclear p-STAT3 (Y705) immunofluorescence (IF) intensity in A549 cells 15min after stimulation with IL-6 (50ng/mL) compared to unstimulated control (un.). Segmentation was based on DAPI-stained nuclei. IF intensity values were normalized to the mean of the unstimulated control (un.) in each experiment and log₂-transformed. Each point represents an individual nucleus (unstimulated = grey, IL-6 = orange); black circles indicate the means from n = 3 independent experiments. Horizontal lines denote replicate means. **C.** Representative Western blot showing total STAT3 (83 kDa) and β-actin (42 kDa) in A549 cells following a 2 h challenge with Spn strains TIGR4 or 6B at MOI 20, or in uninfected controls. **D.** Quantification of STAT3 protein expression in A549 cells after 2-hour challenge with Spn strains TIGR4 and 6B (MOI 20), normalized to uninfected controls (n = 4). Statistical significance was assessed using one-way ANOVA followed by Dunnett’s post-hoc test. ns: p≥0.05.

**Supplementary Figure S2.**
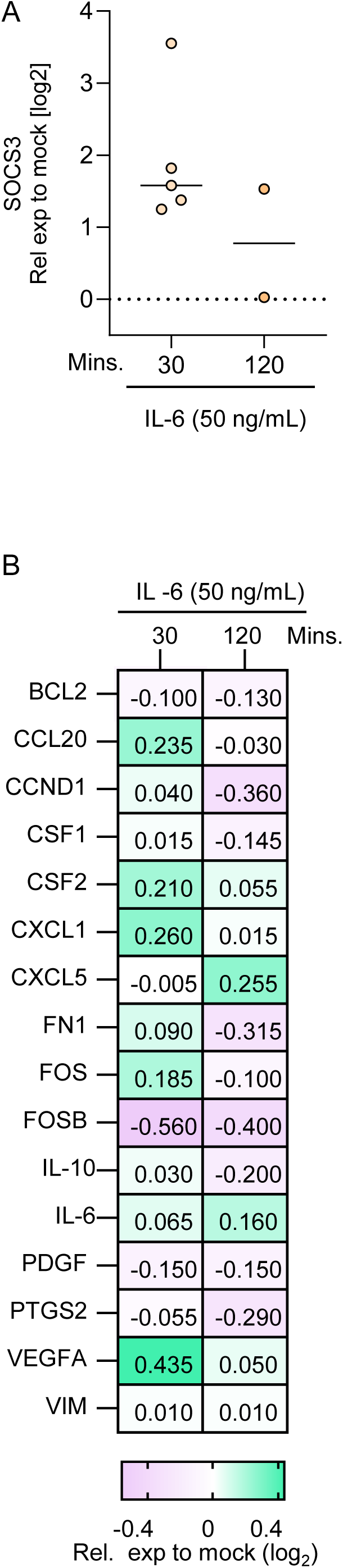
**A.** Semi-quantitative real-time PCR analysis of SOCS3 expression in A549 cells following exposure to IL-6 (50 ng/mL) for 30 min or 2 h. Data are shown as log₂ fold changes relative to the uninfected control from n = 5 (30 min) and n = 2 (2 h) biological replicates. Gene expression was normalized to the housekeeping gene GAPDH and subsequently to the uninfected control (ΔΔCt-method). **B.** Heatmap of real-time PCR results showing expression of selected STAT3-associated genes of interest (GOIs) after 30 min and 2 h exposure to IL-6 in A549 cells. Values represent log₂ fold changes relative to uninfected. The color scale indicates relative expression levels (see scale bar).

**Supplementary Figure S3:**
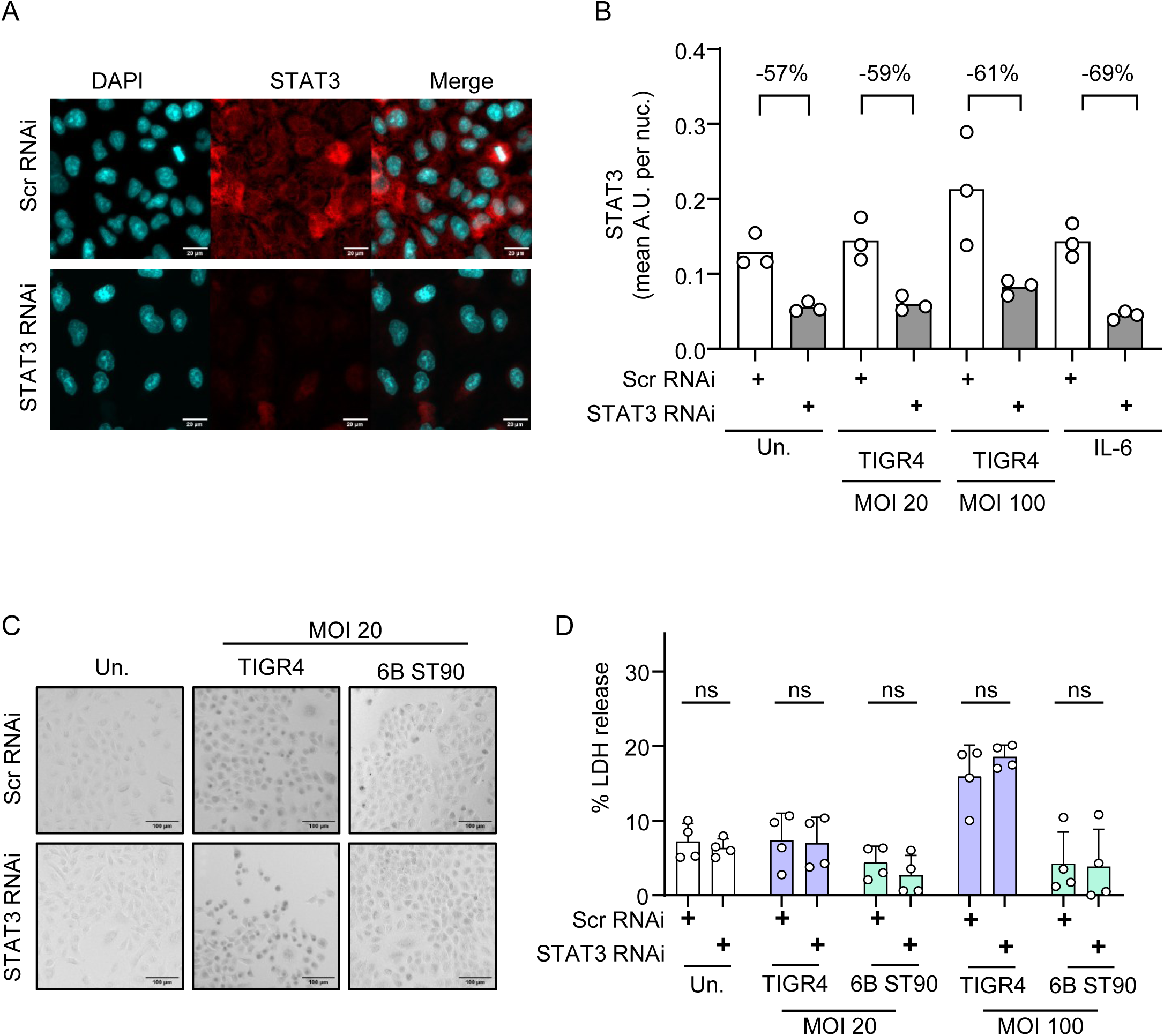
Validation of STAT3 Knockdown Efficiency by RNAi in A549 Cells **A.** Representative immunofluorescence images showing total STAT3 localization in A549 cells treated for 48 h with STAT3-targeting RNAi (STAT3 RNAi) or scrambled control RNAi (scr RNAi). **B.** Quantification of mean total STAT3 fluorescence intensity within the nucleus (segmented by DAPI) in RNAi-treated A549 cells (n = 3 biological replicates) after 48 h of RNAi treatment, followed by either a 2 h challenge with Spn TIGR4 at MOIs 20 or 100, or stimulation with IL-6 (50 ng/mL) for 15 min. Percent knockdown between scr RNAi and STAT3 RNAi is indicated in the figure. **C.** Representative brightfield images (10× magnification) of trypan blue membrane exclusion assay in RNAi-treated A549 cells following 2-hour challenge with Spn TIGR4 and 6B at MOI 20. Cells with compromised membrane integrity show dark blue nuclear staining due to trypan blue uptake. **D.** Lactate dehydrogenase (LDH) release assay in A549 cells after 2-hour challenge with Spn TIGR4 or Spn 6B at MOIs 20 and 100, stimulation with IL-6 (50 ng/mL, 2h), and uninfected controls. LDH release is expressed as a percentage of total cell lysis induced by chemical detergent. Bars represent mean ± SD of n = 4 biological replicates.

**Supplementary Figure S4:**
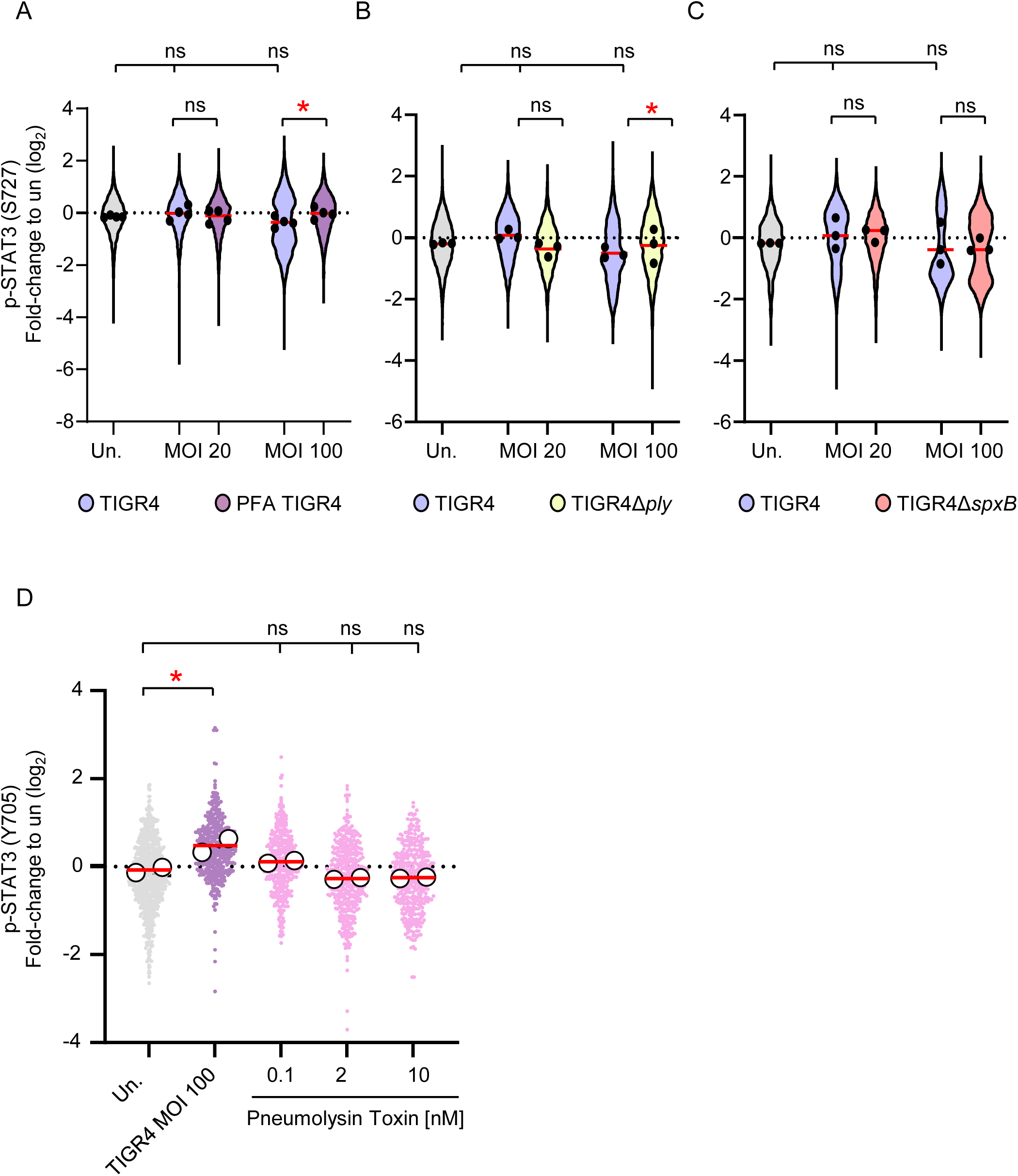
Live *S. pneumoniae* and Pneumolysin Production Are Required to suppress non-canonical nuclear STAT3 localization and recombinant Pneumolysin Alone Is Not Sufficient to Induce Canonical STAT3 Activation in A549 Cells Quantification of phosphorylated STAT3 immunofluorescence intensity (IF) over the nucleus in A549 cells, segmented based on DAPI nuclear staining. Fluorescence intensities were normalized to the mean of the uninfected control(un.) for each experiment and log₂-transformed. Each colored point represents an individual nucleus; black circles indicate the mean of biological replicates, and horizontal lines show the overall mean. **A.** Quantification of p-STAT3 (S727) IF intensity over the nucleus of A549 cells challenged with live Spn TIGR4 and PFA-inactivated TIGR4 (PFA TIGR4) at MOIs of 20 and 100. Color coding: un. (grey), live TIGR4 (purple), inactivated TIGR4 (mauve). (n = 4). Statistical significance was assessed using two-way ANOVA followed by Tukey’s post-hoc test. *p < 0.05; ns, not significant. **B.** Quantification of p-STAT3 (S727) IF intensity over the nucleus of A549 cells challenged with Spn TIGR4 and Spn TIGR4Δ*ply* (pneumolysin-deficient) at MOIs 20 and 100. Color coding: un. (grey), TIGR4 (purple), TIGR4Δ*ply* (yellow). (n = 3). Statistical significance was assessed using two-way ANOVA followed by Tukey’s post-hoc test. *p < 0.05; ns, not significant. **C.** Quantification of p-STAT3 (S727) IF intensity over the nucleus of A549 cells challenged with Spn TIGR4 and Spn TIGR4Δ*spxB* (pyruvate oxidase-deficient) at MOIs 20 and 100. Color coding: un. (grey), TIGR4 (purple), TIGR4Δ*spxB* (salmon). (n = 3). Statistical significance was assessed using two-way ANOVA followed by Tukey’s post-hoc test. *p < 0.05; ns, not significant. **D** Quantification of nuclear p-STAT3 (Y705) IF intensity over the nucleus of A549 cells following a 2 h challenge with Spn TIGR4 (MOI 100) or recombinant pneumolysin (PLY) at 0.1, 2, or 10 nM. Color coding: un., grey; TIGR4, purple; PLY, lilac. (n=2). Statistical analysis was performed using one-way ANOVA followed by Dunnett’s multiple-comparisons test versus uninfected; *p ≤ 0.05; ns, not significant.

## Notes

### Competing Interest Statement

The authors have declared no competing interest.

